# Local and distributed cortical markers of effort expenditure during sustained goal pursuit

**DOI:** 10.1101/2020.12.02.408609

**Authors:** Lauren M. Patrick, Kevin M. Anderson, Avram J. Holmes

## Abstract

The adaptive adjustment of behavior in pursuit of desired goals is critical for survival. To accomplish this complex feat, individuals must weigh the potential benefits of a given course of action against time, energy, and resource costs. Prior research in this domain has greatly advanced understanding of the cortico-striatal circuits that support the anticipation and receipt of desired outcomes, characterizing core aspects of subjective valuation at discrete points in time. However, motivated goal pursuit is not a static or cost neutral process and the brain mechanisms that underlie individual differences in the dynamic updating of effort expenditure across time remain unclear. Here, 38 healthy right-handed participants underwent functional MRI (fMRI) while completing a novel paradigm to examine their willingness to exert physical effort over a prolonged trial, either to obtain monetary rewards or avoid punishments. During sustained goal pursuit, medial prefrontal cortex (mPFC) response scaled with trial-to-trial differences in effort expenditure as a function of both monetary condition and eventual task earnings. Multivariate pattern analysis (MVPA) searchlights were used to examine relations linking prior trial-level effort expenditure to subsequent brain responses to feedback. At reward feedback, whole-brain searchlights identified signals reflecting past effort expenditure in dorsal and ventral mPFC, encompassing broad swaths of frontoparietal and dorsal attention networks. These results suggest a core role for mPFC in scaling effort expenditure during sustained goal pursuit, with the subsequent tracking of effort costs following successful goal attainment extending to incorporate distributed brain networks that support executive functioning and externally oriented attention.

**Significance Statement:** Historically, much of the research on subjective valuation has focused on discrete points in time. Here, we examine brain responses associated with willingness to exert physical effort during the sustained pursuit of desired goals. Our analyses reveal a distributed pattern of brain activity encompassing aspects of ventral mPFC that tracks with trial-level variability in effort expenditure. Indicating that the brain represents echoes of effort at the point of feedback, searchlight analyses revealed signals associated with past effort expenditure in broad swaths of dorsal and medial PFC. These data have important implications for the study of how the brain’s valuation mechanisms contend with the complexity of real-world dynamic environments with relevance for the study of behavior across health and disease.

## Introduction

The successful pursuit of desired goals relies on a set of complex decisions as individuals weigh the potential benefits of a given course of action against the associated costs in terms of time, energy, and resources. While substantial progress has been made studying subcomponents of value-based decision-making, for instance choice and outcome (1, 2), research in this domain in humans has traditionally relied on static approaches that focus on fixed or discrete moments in time. Yet, effortful goal pursuit is a dynamic process and the associated calculations play out in a time-varying manner, likely engaging a diverse array of partially overlapping brain systems. These include cortico-striatal circuits involved in motivated behavior and movement (3) as well as ‘higher-order’ association networks implicated in externally oriented attention (4), the maintenance of task goals, and cognitive control (5–7). Despite the importance of understanding the dynamic functional landscape of sustained goal pursuit, the core features of temporally extended effortful persistence and subsequent responses to task-relevant feedback remain unclear.

The cortico-striatal circuits that underpin subjective value formation (1), choice (8, 9), and goal directed action (10, 11) have been a focused area of study in decision making for many years. In particular, ventral aspects of striatum encompassing the nucleus accumbens and associated swaths of medial prefrontal cortex (1) have been established as core components of the brain’s reward system (3), playing a critical role in intertemporal choice (12), risky and ambiguous decisions (13), as well as reward anticipation and consumption (14–16). These data suggest a critical role for cortico-striatal circuitry in the formation of subjective value (1–17, 18). Complementing this work, tasks examining an individual’s willingness to commit to future effort to obtain rewards have revealed that dorsal aspects of mPFC track positively with effort cost information at cue (19, 20) and during effort prediction error (21). While a role for cortico-striatal circuits in select phases of anticipation and consumption has been established, the contribution of this circuitry in dynamic effortful persistence during active phases of goal pursuit is not as well characterized. Initial evidence suggests that activity in the ventral mPFC tracks the value of continuing to passively persist for an expected reward (22). It is not yet clear whether this mPFC response reflects an individual’s willingness to persist for subsequent reward based on their dynamic and context-sensitive reappraisal of its subjective value or extends beyond passive waiting to track with trial-level variability in physical effort expenditure during dynamic goal pursuit.

A fundamental challenge in uncovering the biological mechanisms that underlie sustained goal pursuit is that even the simplest cognitive tasks rely on an array of cortical and subcortical systems (23, 24), ranging from sensory perception and the generation and implementation of motor plans through subjective value formation and associated adjustments to higher-level task goals (25). Yet, despite this complexity, prior investigations in this domain have largely focused on a curated set of cortico-striatal circuits (26, 27). An alternate but not mutually exclusive possibility is that complex behaviors are instantiated across distributed functional systems throughout the brain. Consistent with this idea, portions of prefrontal cortex thought to support executive functioning and cognitive control, including the dorsal anterior cingulate cortex (dACC), have been found to track choice difficulty (28) and estimated effort costs (29). Additionally, lateral prefrontal cortex territories have been implicated in the extent that future rewards are discounted (30, 31). Broadly, these regions reflect part of a frontoparietal control network (32) that encompasses aspects of dorsolateral and dorsomedial prefrontal, lateral parietal, and posterior temporal cortices as well as corresponding portions of the striatum (33). Yet, while altered intrinsic functional coupling within frontoparietal territories has been observed in patient populations characterized by deficits in hedonic function, including reduced motivation and willingness to expend effort in the pursuit of desired goals (34–37), the extent to which the brain represents the dynamic expenditure of effort across this large-scale network architecture remains to be determined.

In the present study, we use a novel paradigm to investigate brain responses underlying individual variability in participants’ willingness to exert physical effort to obtain monetary rewards or avoid punishments when effort requirements are uncertain. First, examining the active effort phases of our task, we demonstrate a distributed pattern of brain activity encompassing aspects of ventral mPFC that tracks with trial-level variability in effort expenditure as a function of time, monetary incentives, and eventual task earnings. This effect was more pronounced in higher earners and in the reward condition. These results suggest a link between mPFC response and both individual and contextual differences in temporally extended effortful persistence. Next, indicating that the brain represents echoes of trial-level effort at the point of feedback, multivariate searchlight analyses revealed signals associated with past effort expenditure in dorsal and medial PFC. This effort-based response profile was preferentially evident across broad swaths of cortex encompassing both frontoparietal and dorsal attention networks. These collective results suggest a key role for mPFC territories in the scaling of active effort expenditure during goal pursuit. Following successful goal attainment, the subsequent tracking of effort costs is reflected across large-scale brain networks that support executive functioning and externally oriented attention.

## Results

In a sample of 38 neurologically and psychiatrically healthy right-handed participants between the ages of 18 and 35, we used functional magnetic resonance imaging (fMRI) to examine how brain responses track scaling effort expenditure during dynamic goal pursuit. Participants completed a novel Willingness to Exert Physical Effort task (WeePhys; Fig. 1). Over the course of each trial, participants decided how much effort to exert in order to increase the likelihood of either obtaining, or avoiding the loss of, $0.25. To account for individual differences in ability to exert effort, an individual-specific titrated effort value was computed for each participant from their performance during the study practice session. The incentivized portion of the task had 3 reward and 3 punishment blocks with a total of 102 trials. On each trial, participants were instructed to exert as much effort as they would like over 15 seconds by right index finger button pressing. On each trial, a button-press threshold was selected from the individually-specific distribution of required effort (Fig. 1C). If participants met or exceeded this threshold, they earned a reward or avoided monetary loss. If they did not exceed this threshold, they failed to earn a reward or received a monetary penalty depending on the block. On any given trial, participants were not informed of how much effort exertion was necessary to obtain the reward or avoid punishment. To examine the interactions between brain responses and effort expenditure, we first processed the data with a series of steps common to fMRI analyses (see *Methods*).

**Figure 1.**
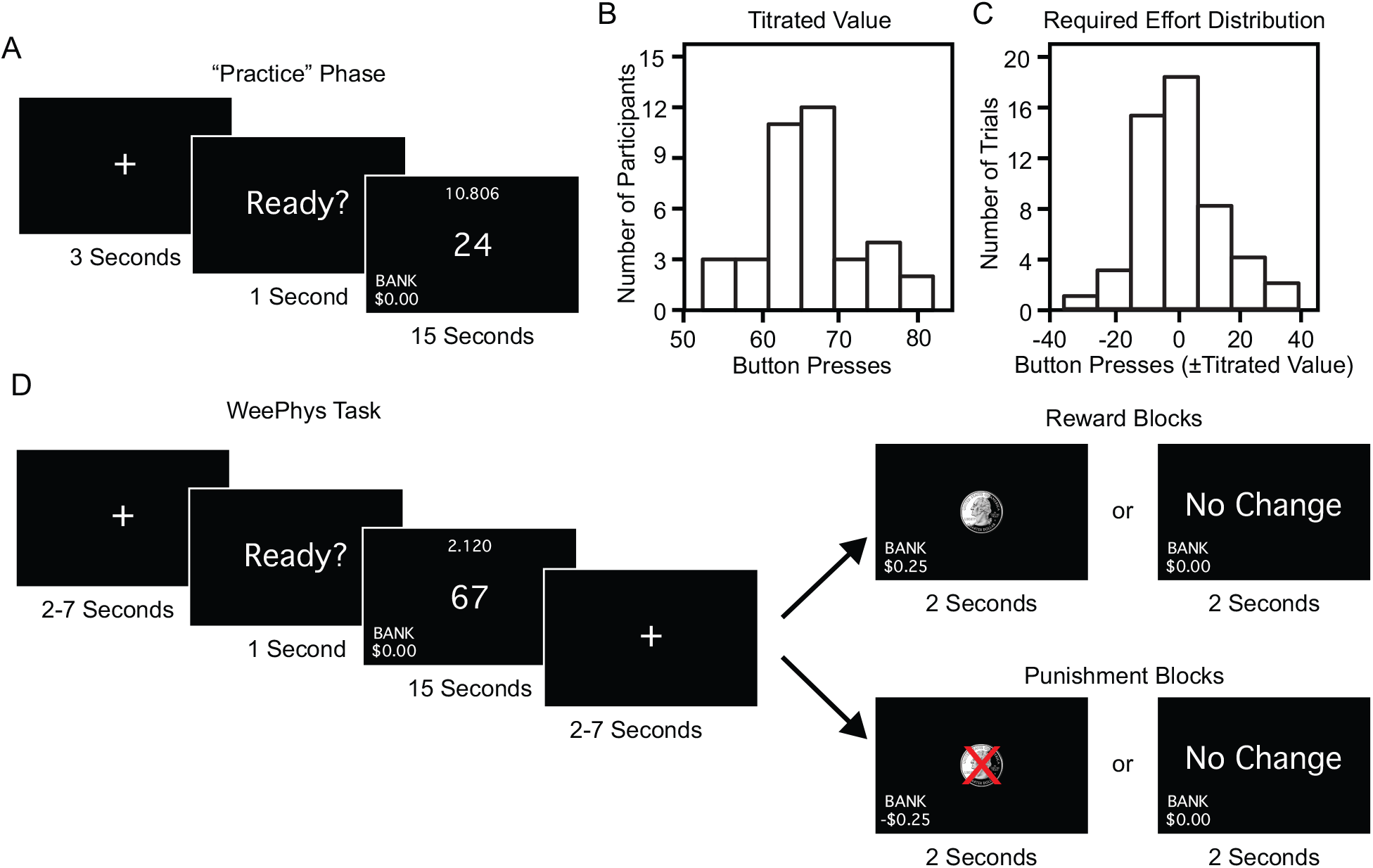
Schematic representation of the Willingness to Exert Physical Effort (WeePhys) task design. (A) Participant specific right index finger button pressing ability is assessed during the practice phase, resulting in a titrated value for each individual. (B) Titrated values are displayed in a histogram reflecting the distribution of button pressing ability across participants. (C) Histogram reflects the distribution of required effort across trials for all participants, centered at each participant specific titrated value. (D) During task performance, participants are successful on a given trial if the number of right index finger key presses exceeds the required number, selected from their individually titrated distribution. Reward and punishment trials are blocked, alternating between runs.

### Intersubject variability in effort expenditure

Behavioral analyses focused on within and between subject differences in effort expenditure. First, consistent with a literature demonstrating heightened performance in incentivized environments (5, 38), we observed a global increase in number of button presses in the task relative to the initial training phase (mean difference from titrated value: 10.61±8.95 (StDev), t(37)=7.31, p≤0.001; Figure 2A). Second, when assessing the expenditure of physical effort, there is the possibility that a portion of the observed variability in participant responses may emerge over time due to accumulating levels of fatigue. Contrary to a pattern that would suggest fatigue, we observed an increase in the amount of effort expenditure between the first (5.43±7.84) and last (12.19±11.43) blocks of the task within participants (t(37)=3.75, p≤0.001; Figure 2B).

**Figure 2.**
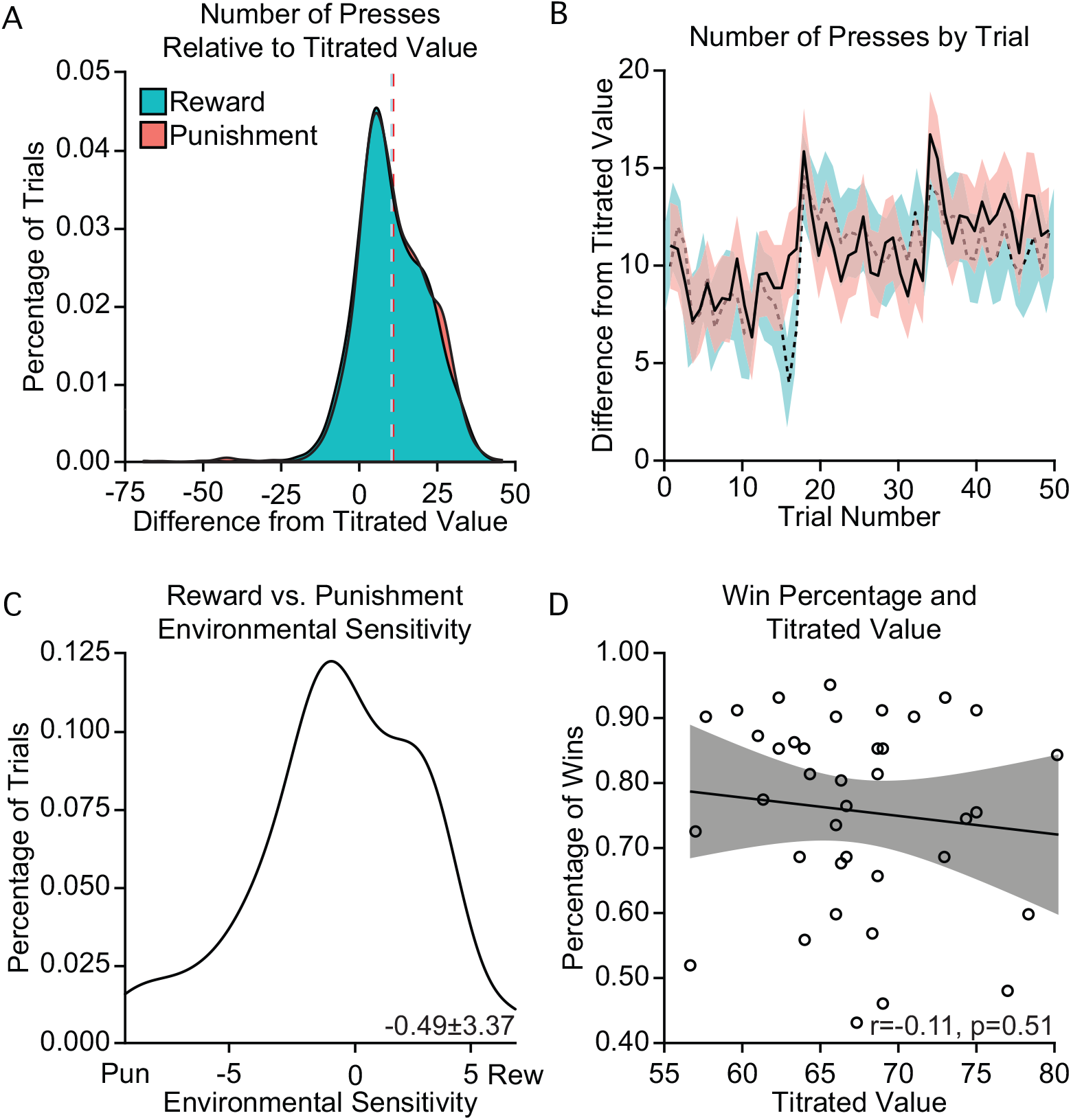
Behavioral task performance. (A) There was a global increase in number of button presses in the task relative to the initial training phase (mean difference from titrated value: 10.61±8.95 (StDev), t(37)=7.31, p≤0.001). (B) Indicating that variability in performance is not driven by fatigue, participants completed a greater average number of button presses over the titrated value on the final task block (12.19±11.43), relative to the first block (5.43±7.84; t(37)=−3.75, p≤0.001). (C) On average, there is no difference in the number of responses compared to the titrated value on reward (10.37±9.01) and punishment trials within participants (10.86±9.20; t(37)=−0.89, p=0.38). (D) Practice task derived titrated values did not track with subsequent task earnings (r(36)=−0.11, p=0.51), suggesting they successfully captured baseline effort expenditure ability without biasing subsequent task performance. Rew, reward trials; Pun, punishment trials. Error bars reflect standard error.

Although individuals are generally more sensitive to potential losses than gains (39, 40), the possible presence of loss aversion during effortful goal pursuit is less clear (41–43). Inconsistent with a loss aversion account of effort expenditure in our data, we did not observe a within-participant difference in effort expenditure between reward (10.37±9.01) and punishment conditions (10.86±9.20; t(37)=−0.89, p=0.38) when compared to the titrated value. Despite no significant difference in conditions at the group level, a subset of participants exhibited nominally increased effort expenditure for punishment relative to reward trials (55%; Figure 2C) indicating a degree of differential sensitivity to reward and punishment blocks across the sample.

It is possible that population-level variability in task performance could, at least in part, depend on the participant-specific distribution of required effort across trials, centered at each individual’s titrated value. Indicating that individual differences in broad button pressing capacity did not drive the observed pattern of results, there was no relationship between the center of the required effort distribution (participant specific titrated value) and task earnings (r(36)=−0.11, p=0.51; Figure 2D). These data suggest that while individual differences in button pressing ability are evident in our sample, they are not driving the observed intersubject variability in goal achievement.

### Medial prefrontal cortex activity tracks trial-level effort expenditure

Functional imaging analyses sought to determine the manner through which brain responses track effort costs during dynamic goal pursuit. Prior work suggests that activity in ventral aspects of mPFC tracks the value of continuing to wait for reward in a given environment (22) but the role of mPFC in scaling effort expenditure in uncertain environments is unclear. To examine the neural correlates of scaling effort expenditure, amplitude modulated signal based on trial level effort expenditure z-scored within subjects was computed. We conducted a whole brain ANCOVA on the effort amplitude modulated signal using AFNI’s 3dMVM (44), examining time (18 seconds from effort onset), condition, and continuous task earnings. When examining the main effect of time, effort expenditure across trials recruited a distributed set of territories encompassing dorsal mPFC as well as swaths of primary motor and visual cortices (Supplemental Figure 1). The observed relationship echoes prior results that have been observed across a diverse set of contexts involving motor activity (38, 45–48), the role of dmPFC in evaluating effortful options (20), and effort learning (21). Although speculative, the observed link between scaling effort expenditure and increased visual system activity may reflect trial-to-trial variability in motivated attention (49, 50) or arousal (51).

Highlighting a role for the mPFC in capturing trial-to-trial level variability in physical effort during dynamic goal pursuit, a significant interaction between time, condition, and total earnings was observed within ventral mPFC (Figure 3; Supplemental Figure 2; p≤0.005, FWE cluster corrected q≤0.01). Post-hoc analyses were conducted to decompose the direction of effects. Follow-up one-way ANOVAs were conducted on the mPFC ROI examining condition (i.e. reward, punishment), overall task earnings and time. First, mPFC showed increased BOLD response for reward relative to punishment blocks. These analyses revealed increased reward block mPFC response (0.07±0.03), relative to the punishment condition during effort expenditure (0.01±0.02; F(1,683)=40.43, p≤0.001). Further, we observed a positive relationship between total participant task earnings and trial-level mPFC response during periods of active effort (F(25,1342)=11.70, p≤0.001; Figure 3). Of note, there was no main effect of time in this portion of mPFC (F(2.9,217.3)=1.76, p=0.16; see Supplemental Figure 1). Together, these results indicate a role for mPFC in successful goal pursuit in contexts where effort requirements are uncertain, varying with task earnings and incentive context.

**Figure 3.**
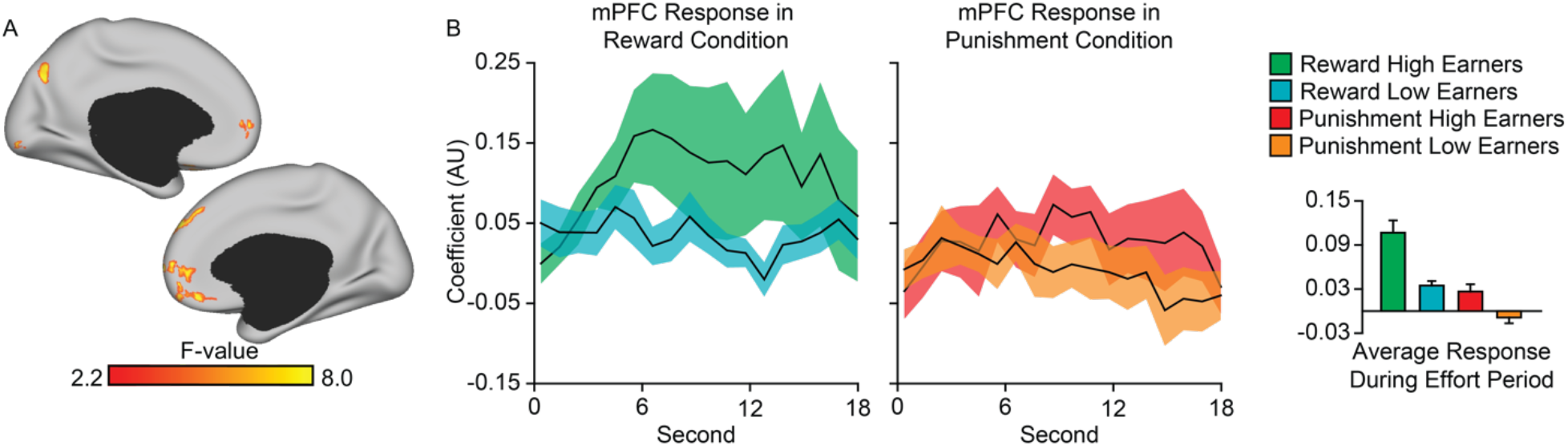
Medial prefrontal cortex (mPFC) activity tracks scaling effort expenditure. (A) Surface map displays ventral aspects of mPFC that track scaling trial-level effort expenditure differentially across feedback condition, earnings and time. Full surface map available in Supplementary Figure 2. (B) The extent that mPFC response scales with trial-to-trial differences in effort expenditure tracks with condition and eventual task earnings over time. Displayed in a median split for ease of visualization, high earners (n=19) show sustained mPFC response, particularly in the reward condition, relative to low earners (n=19). Bar graphs reflect mean mPFC response across the active effort period for reward and punishment trials. Error bars reflect standard error. AU, arbitrary units.

The observed relationships between trial-level effort expenditure and mPFC response echoes prior effects of subjective valuation evident across a broad range of task contexts (1–2, 17). To examine our mPFC effects in the context of this prior work, we quantified the spatial overlap between our whole-brain analyses and canonical valuation-linked circuitry derived through meta-analysis (1). Bilateral clusters within ventral mPFC identified as reflecting the positive effects of value (positive > negative) through meta-analysis and our whole-brain analyses demonstrated an 82-voxel region of overlap in mPFC (5.2% of the canonical region defined through meta-analysis and 23.6% of the empirical cluster). These data suggest that the region of ventral mPFC previously implicated in the encoding of subjective valuation during positive outcomes (1), as well as passive persistence (22), also may reflect the assessment of active effort expenditure during dynamic goal pursuit.

### Prior trial-level effort costs can be decoded at the point of reward feedback

Evidence is accumulating regarding the neural circuits that support the encoding and valuation of physical effort costs (19, 52), however we do not yet know the extent to which sustained effortful persistence is reflected at trial outcome or how these signals differ across environments where there is potential for reward or punishment. To examine the relationship between trial-level effort expenditure and subsequent feedback responses, we swept a 3-voxel radius cubic searchlight using support vector regression (SVR) through the whole brain. For each voxel, we performed two main analyses that examined brain responses to either the successful avoidance of monetary punishment or receipt of reward.

Previous studies have localized effort costs within dorsal mPFC (19, 20), yet sustained goal pursuit is a complex process, relying on attention, control and reward circuitry to dynamically compute the value of options in the environment across changing task contingencies (22, 53). Here, our searchlight analyses were able to successfully decode effort expended over the course of a trial at the point of successful reward feedback (Figure 4; see Supplemental Figure 3 for traditional task contrast of reward and punishment conditions). Consistent with the complex, multifaceted nature of naturalistic goal pursuit, we observed the presence of effort signals distributed across broad aspects of association cortex including territories implicated in cognitive control and externally oriented attention (p≤0.005, FWE cluster corrected q≤0.01). In our data, only searchlight clusters in the reward condition demonstrated significant decoding of trial-level effort expenditure. Consistent with prior work suggesting cortical correlates of gains but not neutral outcomes (54), when examining outcomes indicating successful avoidance of monetary penalty there were no significant relationships linking trial-level effort and subsequent brain responses at feedback. Although speculative, this suggests that while behavioral performance between reward and punishment conditions was similar, the neural circuitry echoing prior effort costs may vary depending on task condition. However, one important caveat to consider when interpreting these results is the low number of unsuccessful trials across both punishment and reward conditions, which prevented us from conducting searchlight analyses on those outcomes. For this reason, we cannot rule out the possibility that brain responses across feedback conditions may broadly reflect prior effort expenditure following periods of both successful and unsuccessful goal pursuit.

**Figure 4.**
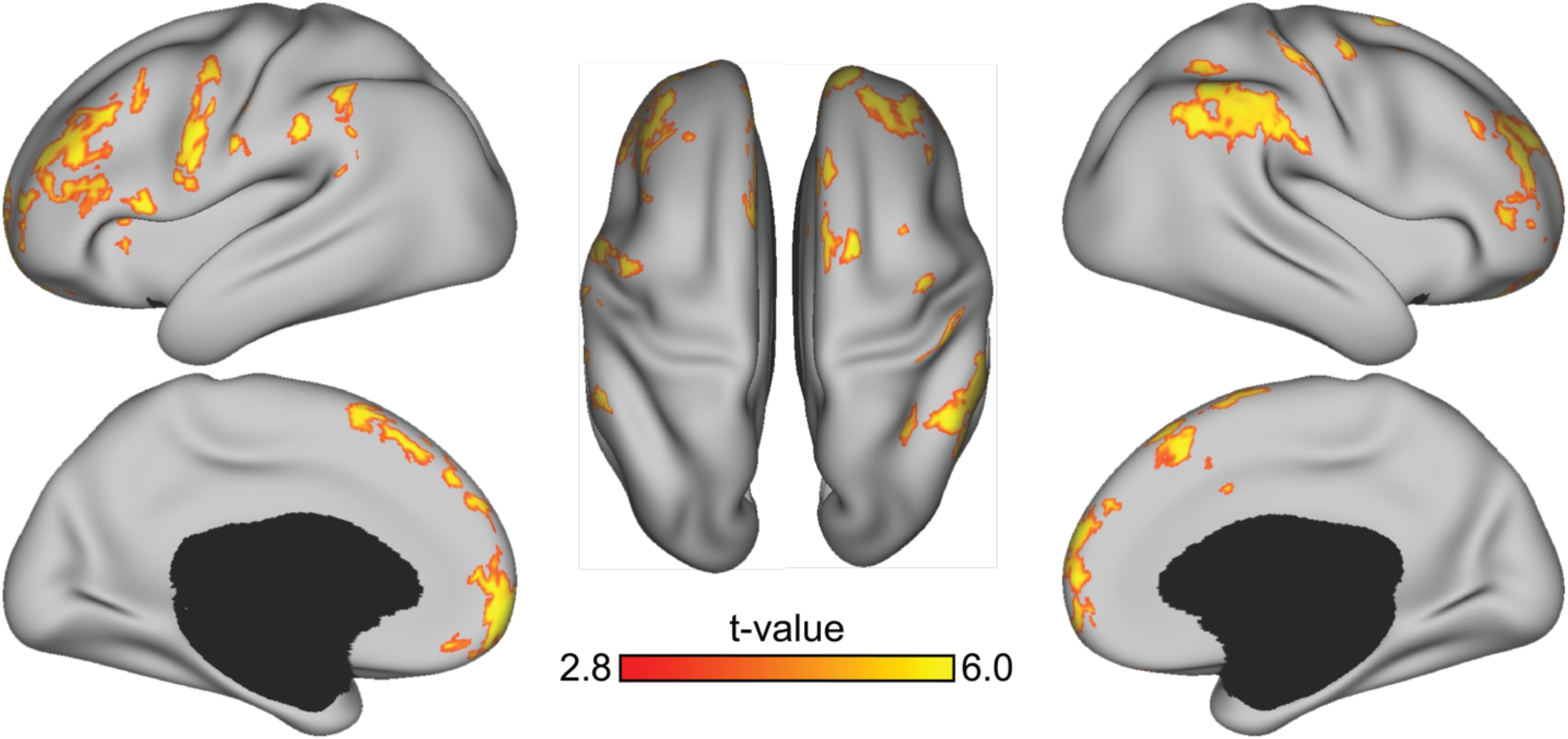
Prior effort expenditure can be decoded at the point of reward feedback. Searchlights using support vector regression (SVR) successfully decoded prior trial-level effort expenditure at the point of subsequent reward feedback in aspects of medial and lateral prefrontal cortices as well as the parietal lobe. Decoding was specific to the reward condition, suggesting that while behavioral performance between reward and punishment conditions was similar, the neural circuitry representing prior effort costs may vary depending on task environment.

### Brain responses tracking prior efforts costs are nonuniformly distributed across cortical networks

The planning and expenditure of physical effort during goal pursuit relies on a diverse set of executive functioning, attention, reward, and motor systems (19, 55), yet we know little about the brain regions that directly underlie dynamic effort computations. To better characterize the spatial distribution of effort decoding across the cortical sheet, we assessed the searchlight accuracy within each of 200 parcels from 7 large-scale brain networks (56) as derived through the cortical parcellation of Schaefer and colleagues (57). Decoding accuracy within the boundary of each network was averaged across parcels and compared (Figure 5c). In the present set of searchlight analyses, the observed effort signals displayed a nonuniform distribution across cortical networks. Decoding accuracy was greatest in heteromodal association cortex including aspects of lateral prefrontal cortex, the temporal-parietal junction, and frontal pole and relatively less pronounced in unimodal sensory and motor as well as orbital frontal cortices. Consistent with this spatial pattern, frontoparietal as well as dorsal and ventral attentional networks demonstrated a high level of decoding accuracy, whereas limbic, somatomotor, and visual systems exhibited the lowest accuracy. Of note, despite encompassing aspects of canonical valuation circuitry including mPFC (1), the default network displayed significantly less decoding accuracy than the frontoparietal network, exhibiting a profile that was consistent with the mean accuracy across cortex (Figure 5c, Supplemental Figure 4). These data indicate a role for networks implicated in executive functioning, cognitive control, and externally oriented attention in integrating effort cost information following periods of active effort expenditure.

**Figure 5.**
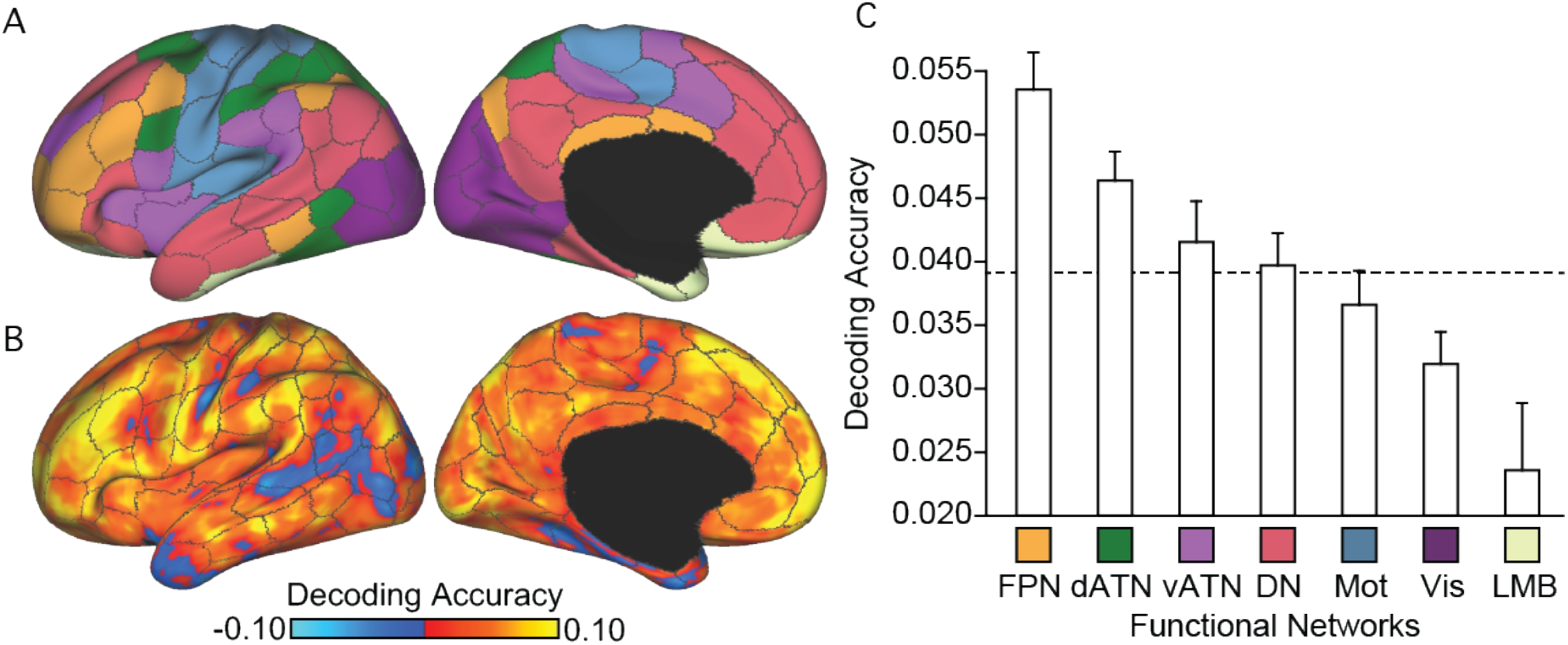
Accuracy of effort decoding is nonuniformly distributed across cortical networks. (A) Analysis were based on the Yeo et al., 2011 7-network solution averaged across the 200 parcel functional atlas of Schaefer and colleagues (2018). (B) Unthresholded decoding maps with the Schaefer parcellation borders overlaid. Scale bar reflects searchlight decoding accuracy for trial-level effort expenditure at the point of reward feedback. (C) Decoding accuracy was highest in frontoparietal and dorsal attention networks. Decoding accuracy within the boundary of each network was averaged and plotted. Error bars reflect standard error. The dotted line indicates the global mean accuracy across the entire cortical sheet. FPN, frontoparietal; dATN dorsal attention; vATN, ventral attention; DN, default; Mot, sensory-motor; Vis, visual; LMB, limbic networks.

## Discussion

Human capacity to weigh the benefits of a potential course of action against the expected effort required to complete it underlies adaptive decision making and foraging behavior in an ever-changing environment. The investigation of reward anticipation and consumption, effort encoding, and their integration over time has long been the subject of empirical investigation in psychology and neuroscience (58–60). However, the bulk of this work examines select phases of goal pursuit and associated responses within a circumscribed ventral cortico-striatal network (3). Here, our study identified that mPFC response scales with trial-to-trial differences in effort expenditure as a function of both monetary condition and eventual task earnings. Providing evidence that brain activity at time of reward consumption tracks exerted effort following successful goal attainment, dorsal and medial prefrontal cortices represented information regarding prior trial-level effort expenditure at the point of reward feedback. Collectively, these observations suggest that effortful persistence emerges from dynamic cost/benefit calculations that are widely distributed across the cortical sheet, preferentially occupying large-scale networks that support executive functioning and stimulus driven attention.

Much is known about the nature of mPFC activity during value-based decision-making (1, 3) and temporal persistence (22), but the part these mechanisms may play in temporally extended periods of effortful goal pursuit remains unclear. Through the use of a continuous behavioral measure, our study identified a clear role for ventral aspects of mPFC in the context of trial-level variability in effortful persistence. In the present task, participants were blind to the required effort necessary on a given trial. Depending on the statistics of a given environment, sustained effortful persistence might increase the probability of a desired outcome, or simply incur increased physical costs without an accompanying benefit. Decision makers work to calibrate their level of effort to minimize costs while maximizing their likelihood of successful goal attainment. Suggesting a role of the mPFC in continuous and adaptive goal pursuit, greater effort expenditure on a given trial was linked with increased BOLD response in ventral mPFC. Providing evidence that this effort related signal links with successful goal pursuit, the extent to which the observed ventral mPFC response tracked with trial-to-trial differences in effort expenditure varied as a function of eventual task earnings, condition, and time. Here, higher earners exhibited greater mPFC response and there was increased mPFC response in the reward relative to punishment condition. These results are consistent with the view that both temporal (22) and dynamic effortful persistence depend on overlapping cortical territories, perhaps sharing cognitive processes with other forms of subjective evaluation and decision making.

Given participants’ continuous motor output throughout the effortful task phase, the present design did not allow for independent examination of effort and motor responses during goal pursuit. As reflected in the ongoing debate regarding the role of frontal midline regions in tracking choice difficulty versus foraging value (28, 61), the role of frontal midline regions in foraging and complex goal pursuit is likely multifold (62). Natural behaviors are often directed towards temporally distant goals that require continuous tracking of targets in the environment. While the local and distributed network-level cortical signals that underlie subcomponents of decision making are still being examined, both tonic and phasic dopamine neurotransmission likely play a critical role in continuous estimates of the proximity of desired rewards, including time judgments (63) and action vigor (64, 65). Tonic dopamine in particular may invigorate action through the coding of average reward (65). However, the extent to which dopamine related processes may play a preferential role in gating effort expenditure remains to be established (66, 67) and it is likely that other neurotransmitters such as serotonin play critical roles as well (68). It will be important for future studies to further examine how *in vivo* measures of effort expenditure in humans can be integrated with bench-lab animal models to better dissociate the molecular and cellular bases of reward and effort signals during goal pursuit (69).

A fundamental question facing the field of decision neuroscience concerns the extent to which goal pursuit, learning, and foraging behaviors are supported through local patterns of activity or are instantiated across the broader large-scale networks of the human brain (70). Tasks that require calculating the expected reward value of a choice (8, 71) have been found to elicit a frontal midline response, while lesions to human ventral aspects of mPFC result in deficits in the representation of the consequences of choices in decision-making tasks (72, 73). Suggesting the role of a broader network architecture supporting ‘higher-order’ executive functions and externally oriented attention in dynamic effortful goal pursuit, our analyses revealed that trial-level effort expenditure can be decoded at the point of reward feedback. The accuracy of this effort decoding was nonuniformly distributed across the cortical sheet, preferentially evident in frontoparietal (32) and dorsal attention networks (4). Regions within the frontoparietal control network are believed to play a crucial role in goal-directed planning (74) the application of complex, nested rules (75), and the monitoring and execution of task sequences in the service of overarching goals (76). Extending upon prior evidence for frontoparietal territories in tracking choice difficulty (28), estimating subsequent effort costs (20, 29) and discounting future rewards (30, 31), the present data suggest a role for frontoparietal network, particularly lateral PFC, in the representation of trial-level effort expenditure at the point of feedback. This finding sheds light on how effort costs may be represented when reward is held constant, something that has been underexamined in the literature (69). Additionally, we were able to decode effort signals on successful reward trials in the absence of explicit information concerning the required trial-level effort, indicating that information about expended effort may be present in the absence of effort prediction error. One interesting future test would be to examine the pre-effort/preparatory phase of goal pursuit to establish the extent to which frontoparietal activity patterns may be predictive of subsequent trial-level effortful persistence.

The present study design is not without limitations. An important future direction will be to utilize tasks that provide robust numbers of trials where participants fail at goal attainment, allowing for the examination of associated brain responses. Given the infrequent occurrence of unsuccessful reward and punishment trials, we are unable to draw conclusions regarding the representation of trial-level effort expenditure for these outcomes in the present analyses. Additionally, as noted above, the use of continuous button pressing during goal pursuit limited our ability to differentiate between striatal motivation, reward, and motor signals during effort expenditure. A related consequence of this design decision is the potential for increased movement on the part of our participants. While the observed results are robust to motion scrubbing and correction, we excluded 9 participants for excessive head movement. Finally, prior work indicates that decision makers adaptively calibrated their level of passive persistence across distinct environmental contexts (22). The extent to which the observed brain responses to ongoing effortful persistence and subsequent feedback are sensitive to global environmental shifts in task demands and/or shifts in outcome probabilities remains an open question.

Select phases of goal pursuit, in particular the anticipation and receipt of desired outcomes, are believed to be supported though activity in mPFC and corresponding aspects of ventral stratum (1–17, 18). A long-standing question concerns how the brain supports the dynamic and continuous updating of effort expenditure throughout motivated goal pursuit. Our analyses revealed a link between ventral mPFC activity and trial-level effortful persistence as a function of time, monetary incentives, and eventual task earnings. Subsequent feedback responses in frontoparietal and dorsal attention networks were found to preferentially reflect prior trial-level effort expenditure. These data suggest the examination of active phases of goal pursuit will yield a more concrete understanding of how the brain’s valuation mechanisms contend with the complexity of real-world dynamic environments with important implications for the study of behavior across health and disease.

## Methods

### Sample Characteristics

Participants between the ages of 18-35 were recruited from the New Haven community (n=53, mean age=23.7±4.9 (StDev), percent female=59). Participants were right-handed and neurologically and psychiatrically healthy, with no structural brain abnormalities. Of these, 15 participants were excluded: 9 for excessive head motion (defined a priori as greater than 10% of frames with greater than 0.3mm motion or maximum displacement >10mm), 2 for abnormally low tSNR (<100), 1 for behavioral evidence of inadequate task performance defined by task earnings below - $4 (on average >15 presses below baseline performance per trial) and 3 for technical difficulties. This yielded datasets from 38 participants (age=24.0±5.1, percent female=63). The reported study procedures were approved by the Yale University Institutional Review Board and written and informed consent was obtained from all participants.

### Procedure

The Willingness to Exert Physical Effort (WeePhys) task was designed to measure how brain response tracks scaling effort expenditure during dynamic goal pursuit (Figure 1). In each task trial, participants decided how much effort to exert in order to increase the likelihood of obtaining $0.25 or avoiding a penalty of $0.25. Maximum earnings on the task were $12.75 and participants were paid their task bonus rounded up to the nearest dollar. Prior to entering the MR scanner, participants completed a “practice” task where on each trial they were instructed to key press on a keyboard as many times as possible over a 15 second window to become acclimated to the task environment. Participants then repeated this 4 trial “practice” sequence in the MRI scanner prior to beginning the incentivized blocks of the task. To account for individual differences in ability to exert effort, the last three practice trials from the in scanner practice block were averaged and used to calculate a titrated effort value for each participant. This titrated value served as the center of the required effort distribution in the incentivized component of the task.

Following the practice phase, participants were introduced to the incentivized version of the task. They were instructed that there would be reward and punishment blocks, as indicated at the beginning of every block, and that on each trial the more effort they exerted, the more likely it would be that they would receive a reward or avoid being punished. However, they were not informed of the amount of effort they would need to expend in a given trial.

In the incentivized component of the task, each participant completed 51 reward and 51 punishment trials (3 blocks reward, 3 blocks punishment, 17 trials per block; Block duration 438s). After a 1s ready period, during the 15s response phase on each trial participants made right index finger key responses. During the effort expenditure portion of each trial, participants were presented with a timer indicating the remaining duration of the trial, a counter reflecting the total number of button presses that had been recorded, and a bank displaying their total earnings in the task. The trial-specific number of responses required to receive reward or avoid punishment was normally distributed across the task (Figure 1c) and centered around an individually titrated baseline number of responses computed during a practice phase of the task (Figure 1b). Jitters of 2-7 seconds were included before the ready period and prior to receiving feedback.

### Image Acquisition

Data were collected on a Siemens Prisma 3T scanner at the Yale University Magnetic Resonance Research Center. Briefly, structural data included a high-resolution, T1-weighted magnetization prepared gradient-echo structural scan (TR=2400ms, TE=2.12ms, 1×1×1mm voxel size). Functional data were acquired using an AC/PC aligned multiband echo-planar pulse sequence (acceleration factor=6, 72 slices, TR=1000ms, TE=33.0ms, 2×2×2mm voxel size, 64-channel head coil).

### Behavioral analysis

Analyses were conducted using R (77). These analyses sought to identify differences between reward and punishment conditions and confirm that fatigue and baseline effort expenditure were not driving observed variability in performance. A one-sample t-test was used to examine differences between baseline effort expenditure (titrated value) and effort expenditure in an incentivized context (overall change in number of responses during the task phase). Paired samples t-tests were used to examine responses on reward and punishment trials, and the number of button presses on the first and last blocks of the task. Pearson correlation was used to examine the relationship between win percentage and titrated value. On average, participants were successful on 76.3% of trials.

### fMRI Analysis

Data were preprocessed using AFNI version 18.2.15 and analyzed using AFNI and Python, including BrainIAK (78), and scikit-learn (79). Functional data were registered to minimum outlier, motion corrected, undistorted and warped to MNI152 space and aligned using the minimum outlier volume and an lpc+zz cost function, outlier-attenuated (AFNI’s 3dDespike), smoothed with a 4mm FWHM Gaussian kernel and intensity-scaled to mean 100 and masked with averaged anatomical volume. The first 5 seconds of each run were removed to allow for T1-equilibration effects.

### Univariate analyses

Voxel-wise general linear models (GLMs) were fit using ordinary least-squares (AFNI’s *3dDeconvolve*). GLMs were estimated for each subject individually using data concatenated across the 6 runs. There were 12 baseline terms per run: a constant, 5 low-frequency drift terms (first-through-fifth-order Legendre polynomials), and 6 motion parameters (roll, pitch, yaw, dS, dL, dP). Event-related BOLD signal time courses were flexibly estimated by fitting piecewise linear splines (‘tent’ basis functions). Amplitude modulated signal based on trial level effort expenditure z-scored within subjects was computed. For trial onset–locked time courses, basis functions were centered every 1s for 28s after stimulus onset. A whole brain ANCOVA-style analysis was implemented on the first 18s after stimulus onset using AFNI’s 3dMVM (44) examining time and condition as within subject variables, earnings as a between subjects variable and centering the order of the timepoints and earnings as quantitative variables in order to lessen any potential correlation between variables in the interaction effects. All whole-brain, group-level analyses were assessed for statistical significance on the basis of FWE cluster correction, with threshold set at p≤0.005, q≤0.01 and cluster estimates computed using 3dClustSim with a mixed-model spatial auto-correlation function (ACF).

### Searchlight analysis

We implemented Support Vector Regression (SVR) within participant using a radial basis function (rbf) kernel (slack parameter C=1 and epsilon 0.1 were chosen a priori) to decode the extent to which feedback responses reflected the amount of trial-specific effort expended during goal pursuit. This portion of the analysis was performed on the timecourse corrected for the 12 baseline terms outlined above. One participant with the fewest successful trials was excluded from this portion of the analysis for having insufficient data for model fit. For each participant, we performed a whole brain 12mm-voxel cubic searchlight analysis (radius=6mm, 3 voxels). Support Vector Machine (SVM) algorithms such as the ones used in this study are the most widely used algorithms for both two-choice classifications as well as regression and are robust and reasonably stable when handling noisy features. Exploring different algorithms would be interesting but could lead to overfitting and would be unlikely to impact performance in a systematic way in the present dataset.

The images used in these analyses were whole-brain activation maps masked to the group. The support vector regression (SVR) model was trained to distinguish signals corresponding to effort expended in that trial for all searchlight cubes including at least 30 voxels. Trials were partitioned into 10 folds and performance was tested on the held-out data from trials not used for training in that fold of cross-validation to minimize overfitting. This was repeated such that each part of the data set was once used for testing (10-fold cross-validation), and the performance was quantified by averaging the fisher z-transformed correlation coefficient reflecting the relationship between actual and predicted effort expenditure in the test data across the folds. This correlation coefficient was then assigned to the center voxel of each searchlight. Intersubject reliability was assessed by applying a one-sample t-test (against zero, using 3dttest++) of the decoding accuracies for each voxel. Group-level analyses assessed for statistical significance on the basis of cluster-based FWE correction, with threshold set at q≤0.01.

### Network analyses of effort decoding accuracy at time of reward

To characterize the spatial distribution of effort decoding across the cortical sheet, the searchlight accuracy at the time of reward was also assessed within each of 200 roughly symmetric volumetric ROIs from 7 specific brain networks (56) in the left and right hemispheres as derived through the cortical parcellation of Schaefer and colleagues (57). Decoding accuracy was first averaged within parcels and then by network (Figure 5c).

## Acknowledgements

This work was supported by the National Institute of Mental Health (Grant R01MH120080 to A.J.H.), the National Science Foundation (DGE-1122492 to K.M.A.), and the Yale Imaging Fund. Analyses were made possible by the high-performance computing facilities provided through the Yale Center for Research Computing. We would like to thank David Gruskin for assistance with data collection and Holmes Lab members for comments on early versions of this manuscript.

---

**Supplemental Figure 1.**
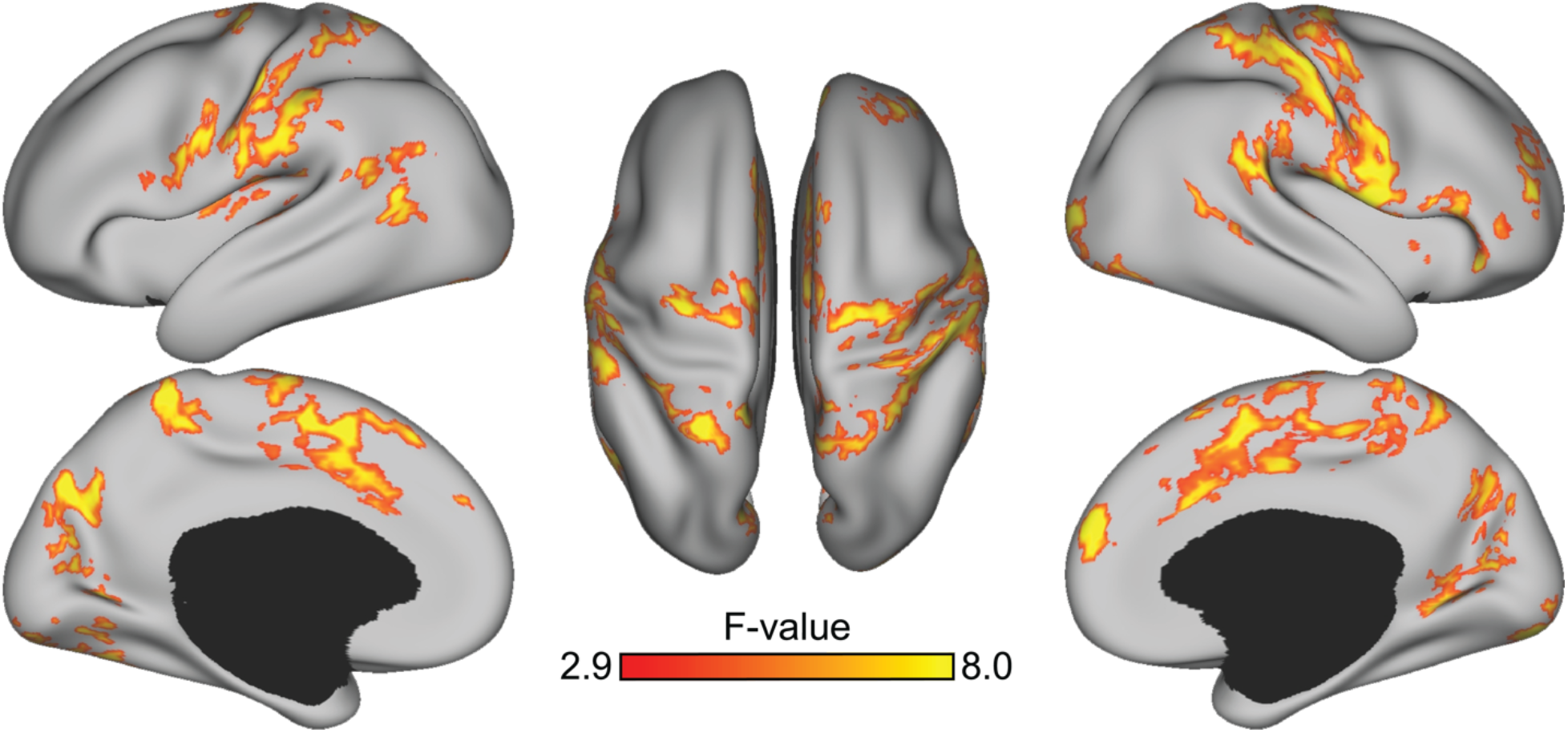
A distributed pattern of brain activity scales with trial-level effort expenditure when examining the main effect of time. Within participants, increased effort expenditure on a trial was linked with greater activity in dorsal aspects of mPFC, along with motor and visual territories.

**Supplemental Figure 2.**
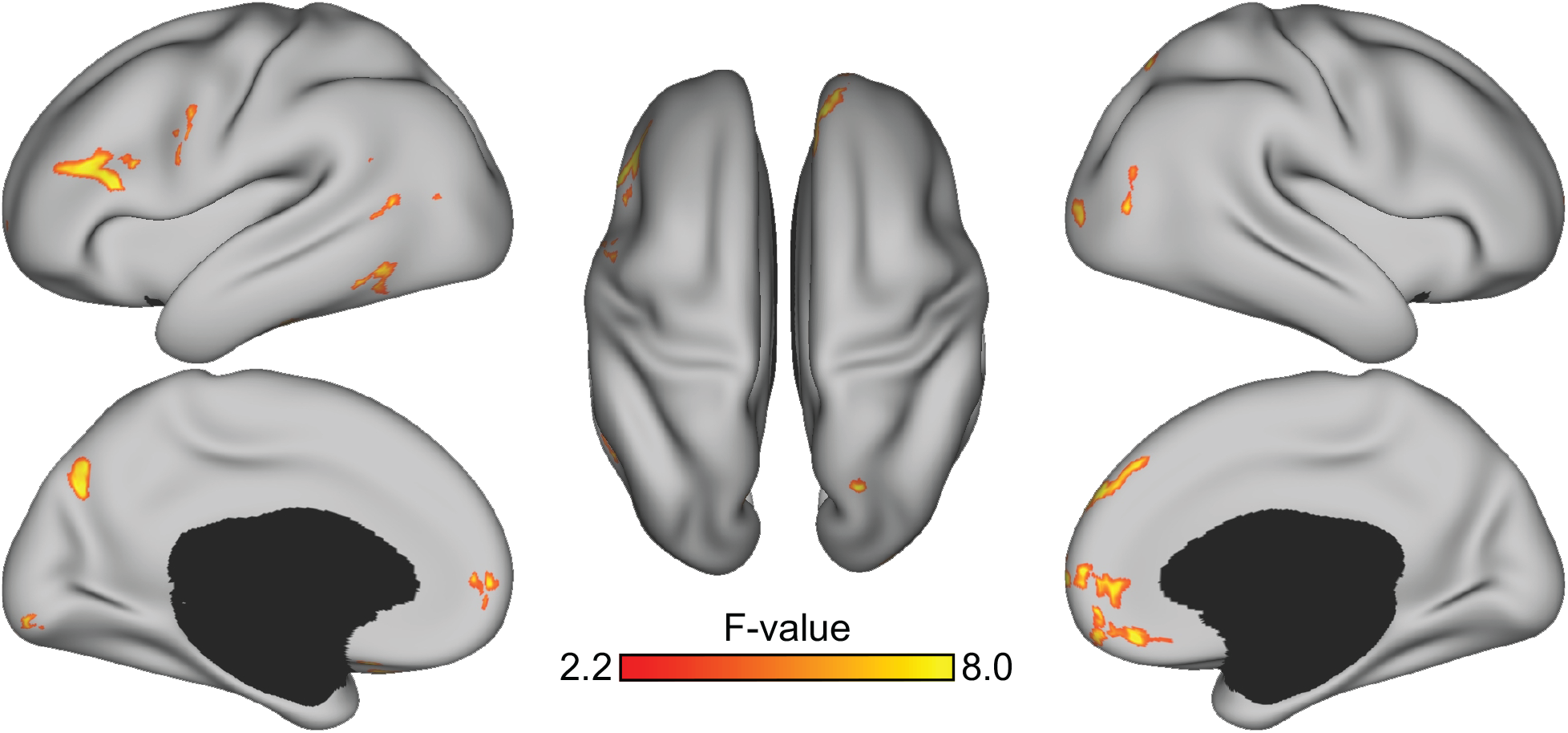
A distributed pattern of brain activity scales with trial-level effort expenditure differentially across feedback condition, earnings and time. Within participants, an interaction between condition (reward or punishment), task earnings, and timepoint in the trial tracked with activity in ventral aspects of mPFC as well as dorsal medial PFC in the right hemisphere and dorsolateral PFC in the left hemisphere.

**Supplemental Figure 3.**
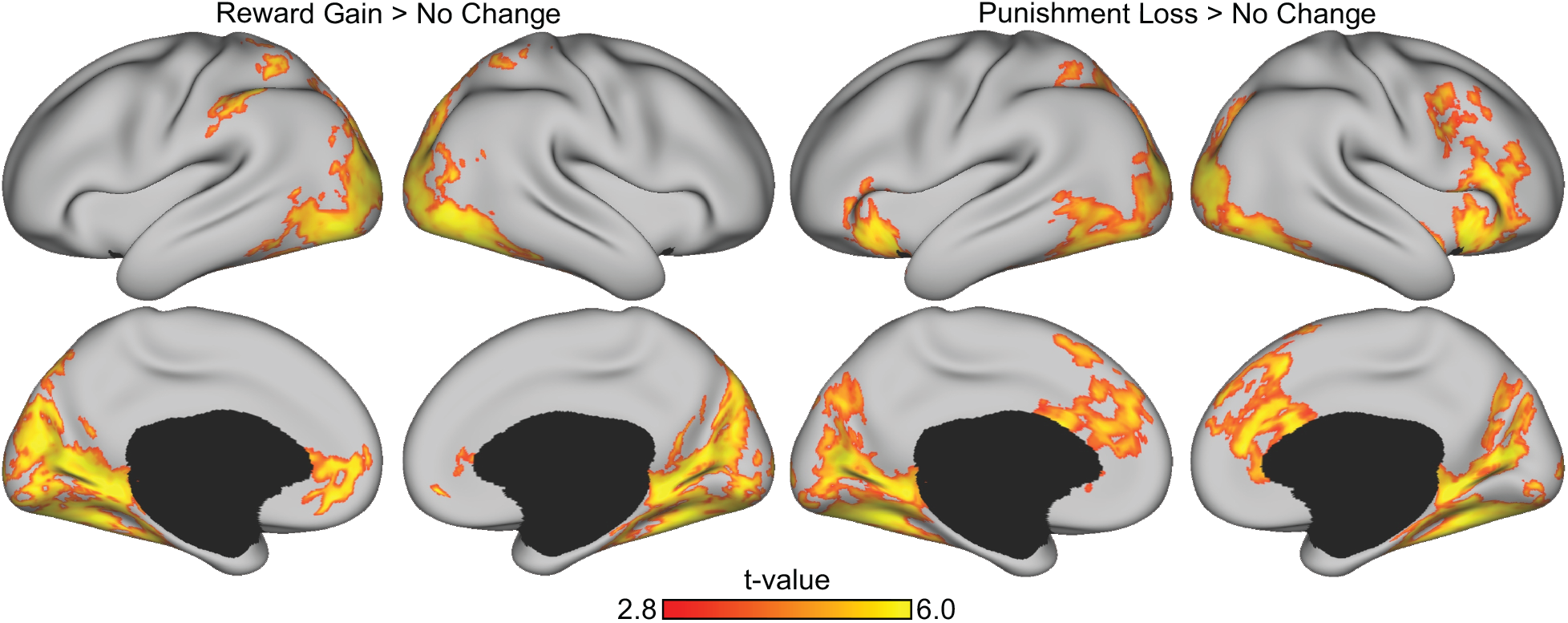
Reward and punishment conditions recruit canonical circuitry. (A) Reward receipt compared to the no change condition preferentially recruited ventral but not dorsal aspects of mPFC. (B) Conversely, punishment receipt compared to the no change condition recruited dorsal mPFC and not ventral mPFC.

**Supplemental Figure 4.**
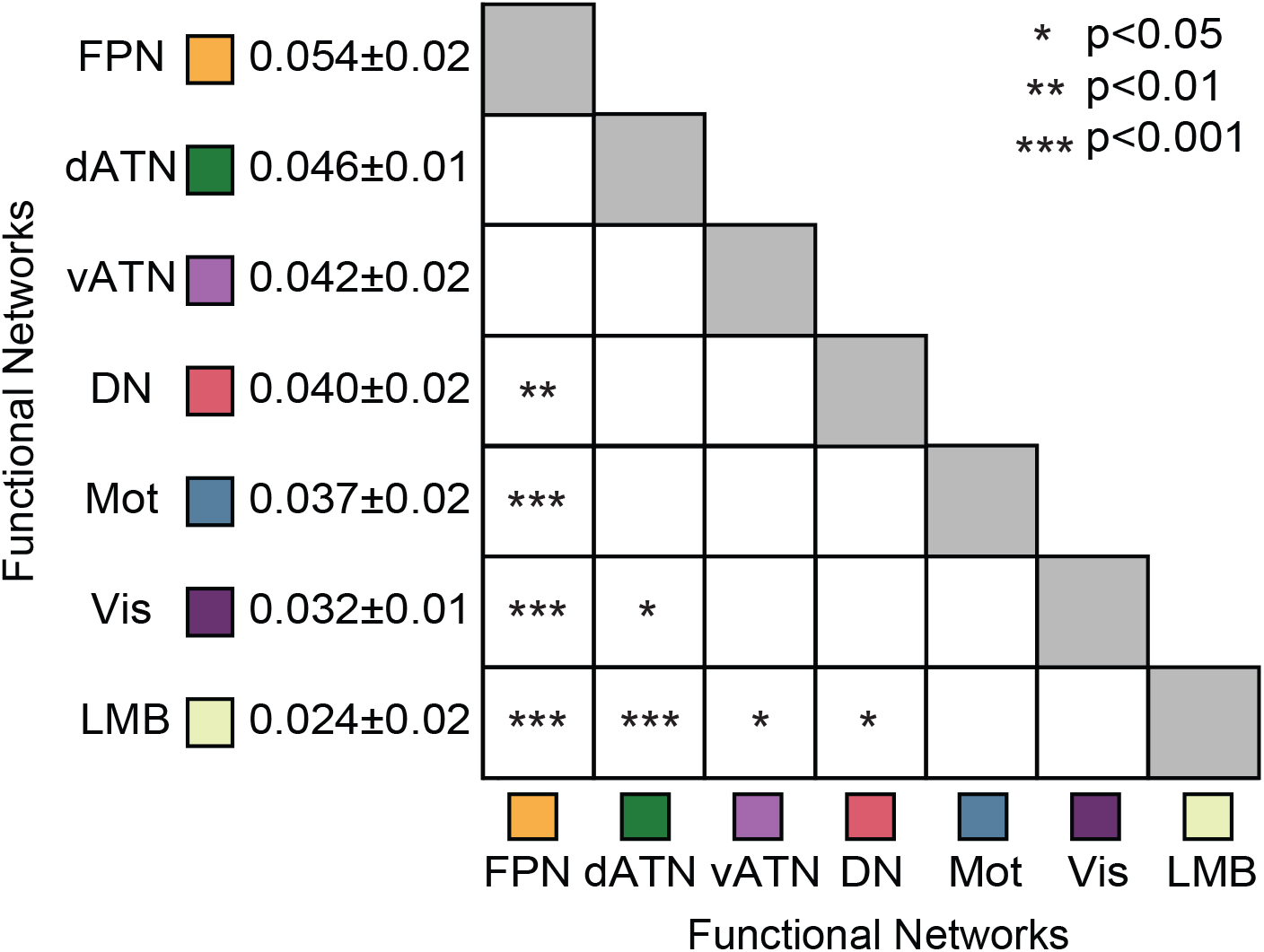
Accuracy of effort decoding was preferential to frontoparietal and attention networks. Analyses were based on the Yeo et al., 2011 7-network solution averaged across the 200 parcel functional atlas of Schaefer and colleagues (2018). P-values Tukey post-hoc corrected. FPN, frontoparietal; dATN, dorsal attention; vATN, ventral attention; DN, default; Mot, sensory-motor; Vis, visual; LMB, limbic networks. Numbers reflect mean ± Standard Deviation.

